# Joint embedding of biological networks for cross-species functional alignment

**DOI:** 10.1101/2022.01.17.476697

**Authors:** Lechuan Li, Ruth Dannenfelser, Yu Zhu, Nathaniel Hejduk, Santiago Segarra, Vicky Yao

## Abstract

Model organisms are widely used to better understand the molecular causes of human disease. While sequence similarity greatly aids this transfer, sequence similarity does not imply functional similarity, and thus, several current approaches incorporate protein-protein interactions (PPIs) to help map findings between species. Existing transfer methods either formulate the alignment problem as a matching problem which pits network features against known orthology, or more recently, as a joint embedding problem. Here, we propose a novel state-of-the-art joint embedding solution: Embeddings to Network Alignment (ETNA). More specifically, ETNA generates individual network embeddings based on network topological structures and then uses a Natural Language Processing-inspired cross-training approach to align the two embeddings using sequence orthologs. The final embedding preserves both within and between species gene functional relationships, and we demonstrate that it captures both pairwise and group functional relevance. In addition, ETNA’s embeddings can be used to transfer genetic interactions across species and identify phenotypic alignments, laying the groundwork for potential opportunities for drug repurposing and translational studies.

## 1 Introduction

Many critical discoveries in medicine have been fueled by a better understanding of molecular mechanisms in model organisms, and the importance of leveraging these models for translational studies continues to increase [1]. However, one of the major challenges to realizing the full potential of model organism studies is functional knowledge transfer [2], the process by which information learned in one species is applied to better understand function in a different species.

Model organisms are widely used to study fundamental biological pathways and disease etiology, given the technical and ethical limitations of performing direct research on humans [3]. Organism complexity inherently has created an imbalance in existing knowledge, as some assays and techniques can be well characterized in simpler organisms such as synthetic lethality studies in yeast [4], and genetic screens in organisms with short life cycles (e.g., worm and fly) [5, 6]. This also provides unique opportunities to capture different biological processes and pathways that might be useful for a researcher’s organism of interest. The ability to transfer meaningful molecular insights from one species to another is therefore a central problem encountered by many experimental biologists, that can both help reveal the broader implications of their own current research as well as guide further experimentation.

Proteins do not carry out their functional roles in isolation, but instead work together, forming biological pathways. Protein-protein interaction (PPI) networks summarize one aspect: physical interactions between protein pairs. For this reason, simply assigning conserved function between species through orthologous proteins is insufficient; furthermore, in many organisms, similar functions can be taken on by proteins that may not be the most sequence similar, but instead take on similar roles in a biological pathway [2]. Thus, one critical component of functional information transfer is the simultaneous “alignment” of biological networks such as PPI networks together with orthology information. Many such network alignment methods have been developed (Table 1). For example, there are methods that have leveraged PageRank [7, 8], genetic algorithms [9], or search algorithms [10, 11, 12], as well as hub alignment-[13] and graphlet-based methods [14]. However, these existing alignment methods usually optimize a convex combination of sequence similarity and topological features.

**Table 1.** Comparison of ETNA with existing unsupervised pairwise network alignment methods.

| method | interpretable embeddings | cross-species anchors | combining sequence & topology | directionality | algorithm | code availability |
| --- | --- | --- | --- | --- | --- | --- |
| **ETNA** | **yes** | **orthologs** | **non-linearly** | **bidirectional** | **autoencoders** | **python** |
| MUNK [15] | yes | orthologs | non-linearly | directional | SVD | python |
| IsoRank [7] | no | BLAST | linearly | bidirectional | PageRank | linux executable |
| HubAlign [13] | no | BLAST | linearly | bidirectional* | minimum-degree heuristic algorithm | C++ |
| PrimAlign [8] | no | BLAST | linearly | bidirectional | Markov chain + PageRank | not available |
| SANA [12] | no | BLAST | linearly | bidirectional* | simulated annealing | webserver, C++ |
| L-GRAAL [14] | no | BLAST | linearly | bidirectional | Lagrangian graphlet | C++, not working |
| GHOST [11] | no | GO | does not use sequence | bidirectional* | seed-and-extend with local search | C++ |
| NETAL [10] | no | GO | does not use sequence | bidirectional* | greedy search on similarity matrix | webserver |
| MAGNA++ [9] | no | n/a | does not use sequence | bidirectional | genetic algorithm | executable, C++ |

\* The method has a bidirectional objective function, but the returned output is directional.

Formulating this alignment problem as a matching problem has limitations [15, 16]—one main limitation is that network topology and sequence similarity are, in some sense, pitted against each other. A recent method, MUNK, seeks to address these limitations by reframing the alignment problem as a joint embedding problem [15]. Specifically, it uses a regularized Laplacian kernel to embed each PPI network individually and uses orthology to project one embedding to the other, achieving better performance than traditional biological network alignment methods for functional tasks. However, MUNK’s embeddings are directional, requiring a decision between ‘source’ and ‘target’ species, leading to differing results for the alignment between the same pair of species. In addition, the latent embedding it uses is the same dimensionality as the smaller of the two PPI networks, which does not fully take advantage of the natural “compression” that is inherent to embedding methods and makes it prone to overfitting.

In the machine learning community, there have been several interesting network embedding methods developed: random walk-based methods [17, 18], matrix factorization-based methods [19, 20, 21, 22], and autoencoder-based methods [23, 24]. Random walk-based methods use random walks combined with skip-gram to learn a node embedding for a network and are capable of capturing information from long-range neighbors. Recently, a matrix factorization-based method called NetMF has been proposed that unifies random walk-based methods into a closed-form matrix that can be solved using singular value decomposition [19]. NetMF’s closed-form matrix contains global network topology information, but since it relies on SVD, it cannot sufficiently capture the complex non-linear relationships inherent in network topology. SDNE is an autoencoder-based method that excels at learning the highly non-linear structure in the network [23]. However, it only considers direct and 2-hop neighbors (i.e., ‘friends of friends’) and ignores longer-range network topology.

Here, we present Embedding to Network Alignment (ETNA), a deep learning method for estimating functional relevance between genes from different species (Figure 1). ETNA leverages the strengths of previous methods in an autoencoder architecture to create individual network embeddings that preserve global and local topological structures of biological networks. Taking inspiration from advances in Natural Language Processing (NLP) by swapping encoders and decoders in a cross-training framework [25] and using orthologous genes as anchors, ETNA aligns two embeddings to a joint latent space. The alignment process allows information from the species to refine individual network embeddings. The final output of ETNA is a bidirectional joint embedding that encodes both within and between species functional relationships.

**Figure 1.**
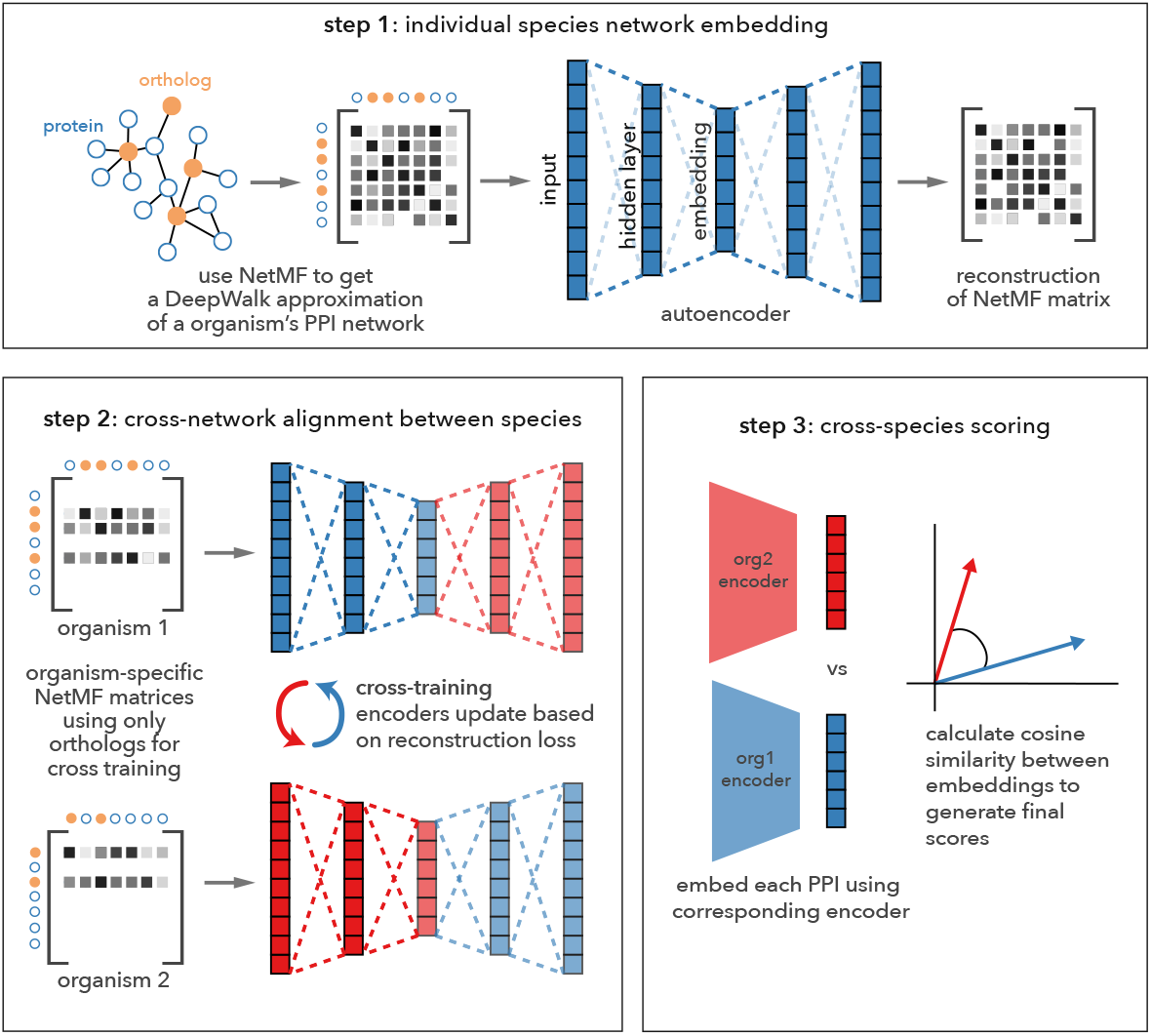
Overview of the ETNA method. ETNA’s framework is roughly divided into 3 main parts: (1) training an autoencoder to embed PPI networks, (2) aligning the embeddings between species using orthologous proteins as anchors for cross-training, and (3) scoring pairs of genes across organisms based on their cosine similarities in embedding space. The embeddings and cosine similarities can both be used for downstream tasks to predict functional similarity across species.

Using human and four other model organisms, we show that ETNA’s joint embedding can capture both pairwise and group functional relationships across species better than the three existing network alignment methods considered as benchmarks. We further explore applications of the joint embedding, including predicting genetic interactions, identifying potential phenotypic alignments between human and mouse, as well as providing new insights on relationships between human disease, mouse phenotypes, and drug targets.

## 2 Methods

### 2.1 A method for cross-species network alignment

To align two networks, ETNA iterates between two main steps: (1) calculating an embedding for each individual network; and (2) aligning the two embedding spaces using orthologs as reference anchors (steps 1 and 2 in Figure 1). By doing so, ETNA identifies a joint embedding between two networks. When applied to PPI networks, this joint embedding can be interpreted as a latent functional similarity map and enables reasoning about relationships between proteins across organisms, beyond ortholog pairs.

#### 2.1.1 Individual network embedding

ETNA uses an autoencoder framework to generate lower-dimensional latent embeddings that preserve both local and global network topology while capturing the non-linear relationships in the input network. More precisely, we modify and extend the SDNE method [23] to incorporate higher-order proximity [26] information beyond second-order. The framework is agnostic to the type of network, but here we focus on PPI networks. Specifically, given a PPI network, we represent it as an undirected graph *G* = (*V, E*), where *V* = {*v*_1_, *v*_2_, · · ·, *v*_*n*_} is the set of *n* proteins and *E* is the set of reported physical interactions between pairs of proteins.

Importantly, instead of using *G*’s adjacency matrix *A* directly as input to the autoencoder, ETNA uses a closed-form approximation of the random walk process on *G*, which we denote by *M* (Equation (3)). The autoencoder is composed of two parts, namely an encoder and a decoder. For each vertex *v*_*i*_, the encoder compresses the *n*-dimensional input (i.e., the *i*th row of *M*, denoted by *m*_*i*_ in the following equations) through a 1024-dimensional hidden layer to a 128-dimensional latent embedding *z*_*i*_. We collect vectors *z*_*i*_ for different vertices into a matrix *Z* ∈ ℝ^*n*×128^ as its rows. The decoder then uses an independent 1024-dimensional hidden layer to map the latent embedding back into reconstruction space (Figure 1). To capture local and global network structure, ETNA uses the following objective function:

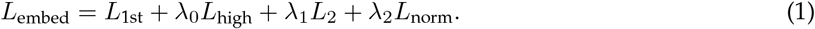

The *L*_1st_ loss preserves the local first-order structure of the PPI network by maximizing the similarity of embeddings between vertices that are directly connected. On the other hand, *L*_high_ captures global network topology by modifying the standard autoencoder reconstruction loss to encourage proteins that have similar network relationships to have similar embeddings. These two losses are described in more detail below. Also, we consider two regularization terms: *L*_2_ norm on the autoencoder parameters to avoid overfitting and 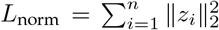 to avoid exploding norms. All hyperparameters are tuned by cross validation (Section 2.3).

##### Preserving local first-order structure

In most real-world networks, the presence of a shared edge between two vertices is a strong signal of similarity [27], and this is true in PPI networks as well. Two proteins that physically interact with each other are more likely to be performing similar functions. The *first-order proximity* captures this local pairwise structure between two vertices. In ETNA, for a pair of vertices (*v*_*i*_, *v*_*j*_) connected by an edge, their latent embeddings should be similar. Thus, the objective function for *first-order proximity* is defined as:

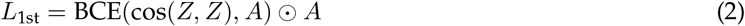

where BCE is the binary cross entropy and ⊙ is the Hadamard product. The term cos(*Z, Z*) ∈ ℝ^*n*×*n*^ denotes a matrix whose (*i, j*)th entry equals 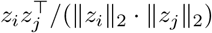 and reflects the similarity between *v*_*i*_ and *v*_*j*_ in the latent embedding space. The Hadamard product with *A* is used here to ensure that only the BCE of known PPIs contribute to *L*_1st_. There are two important reasons for this: i) Physical interactions are not the only criteria to determine whether two proteins are functionally similar, and ii) PPI network data are not yet complete and definitely have false negatives.

##### Preserving global structure through higher-order proximity

Considering only direct neighbor relationships is likely unable to fully capture more complex relationships in a network (e.g., complex pathways), which can involve larger network connectivity patterns. In the past decade, random walk-based methods such as DeepWalk [17] have been shown to learn latent representations that successfully capture network topology. Recently, the NetMF method has been proposed based on a closed-form estimate of the similarity matrix that is implicitly factorized in DeepWalk [19]. Compared to the adjacency matrix, the NetMF matrix not only contains information for direct connections, but also contains similarity values between vertices that are not directly connected. Using its rows as input enables ETNA to consider the higher-order proximity of the PPI network. The NetMF matrix *M* ∈ ℝ^*n*×*n*^ is calculated from the adjacency matrix *A* as [19]:

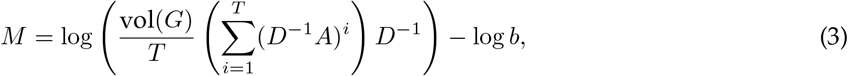

where *D* is the diagonal degree matrix whose entry *D*_*i,i*_ is *v*_*i*_’s degree, 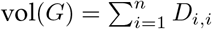 is the volume of graph *G*, *T* is the context window size, and *b* is the negative sampling parameter. Intuitively, *M*_*i,j*_ can be considered as a weighted count of the number of paths from *v*_*i*_ to *v*_*j*_ that have lengths no greater than *T*. ETNA uses the rows of *M* as input into its autoencoder and estimates a reconstruction *M* from the latent embedding. Good reconstructions imply that *higher-order proximity* has been well-captured in the latent embedding, and thus the objective function is defined as:

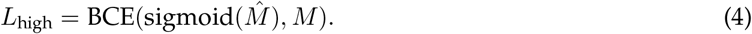

#### 2.1.2 Using orthologs to anchor alignment across species

To align the two previously independent embeddings, we use orthologous proteins as “anchors” between two different species. The underlying intuition is to encourage orthologous protein pairs from the two species to have similar latent features, all while keeping the distance relationship between vertices within each network. Instead of arbitrarily assigning source and target networks and having a directional projection, we use a cross-training method that pushes the embeddings of both networks towards a joint latent space simultaneously. The final embedding in the joint latent space thus contains distance relationships between proteins across networks.

Specifically, the alignment process uses a cross-training method inspired from language translation methods developed in NLP [25]. Recall that we defined a PPI network as an undirected graph *G* = (*V, E*), where |*V*| = *n*. Now, given a second PPI network *G′* = (*V′*, *E′*) with |*V′*| = *n′* and a set of orthologous pairs Θ ⊂ *V* × *V′*, consider an orthologous pair (*θ*_*i*_, *θ*_*i′*_) ∈ Θ: if *θ*_*i*_ and *θ*_*i′*_ play similar roles in their respective networks (i.e., the same “word” in different languages), then we seek to identify a joint embedding (i.e., latent semantic space in NLP or latent functional space for PPIs). If such a joint embedding exists, then one way of thinking about the “translation” task is that once *G*’s encoder places *θ*_*i*_ in the joint embedding, then *G′*’s decoder should be able to reconstruct *θ*_*i′*_’s neighborhood structure, and vice versa. This intuition leads to the following objective function:

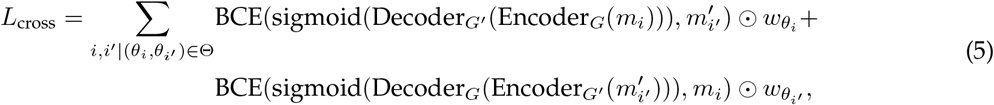

where {*m*_1_, *..., m*_*n*_} and 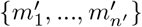 are the row vectors of the corresponding NetMF matrices of *G* and *G′*, and 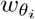 and 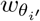 are weights that downscale the contribution of hub genes in the alignment process. Indeed, there are some genes that have multiple orthologous matchings in the other species due to either study bias or their functionality. To make the alignment process not focus only on those hub genes, we set the weight parameters *w* as the inverse of the number of occurrences of a gene in all orthologous pairs.

The alignment process only updates the weights in the encoders, since the alignment should happen in the latent space rather than the reconstruction space. The training process iterates between the embedding and alignment steps to generate a joint embedding that preserves both within- and between-species closeness. The reasoning here is that well-trained individual network embeddings are too “rigid” to accommodate between-species protein closeness. Therefore, a tunable number of embedding process epochs followed by one epoch of alignment process is considered as one training block for ETNA, and the number of training block epochs is decided by cross-validation (Section 2.3). After training the method, the final similarity between any two genes is represented by the cosine similarity between their embedded features in the joint latent space. More precisely, we compute a score matrix *S* = cos(*Z, Z′*) ∈ ℝ^*n*×*n′*^, which contains pairwise similarities between all vertices in *G* and *G′* (step 3 in Figure 1).

### 2.2 PPI network and orthology data

PPI data for each of the six species studied in this work were downloaded from BioGRID (v3.5.187) [28]. Using Entrez gene identifiers, we constructed an unweighted and undirected species-specific PPI network, filtering out self-loops. We also recursively filtered out vertices with the same neighborhood structure, since with only topological information as input, these are indistinguishable to our model. Orthology data were downloaded from OrthoMCL (v6.1) [29].

### 2.3 Building a cross-species functional evaluation standard

The Gene Ontology (GO) provides valuable gene function annotations across species, where genes annotated to the same terms can be considered as functionally similar. We used GO (2020-07-16) [30] as the gold standard to evaluate the ability of our embedding-alignment method to capture cross-species functional similarity. We restricted annotations to the Biological Process (BP) aspect, which describes the molecular activities of genes, and used low throughput experimental evidence codes (EXP, IDA, IMP, IGI, IEP), excluding evidence code IPI (Inferred from Physical Interaction) to avoid introducing any circularity to the evaluations. All GO annotations were propagated through the “is a” and “part of” relations. We restricted our set of GO terms to a slim set representing specific diverse functions that are present across human and four major model organisms (*Mus musculus*, *Saccharomyces cerevisiae*, *Caenorhabditis elegans*, and *Drosophila melanogaster*). We defined specificity as terms with at least 10 genes and at most 100 genes annotated. We also included previously expert-curated GO slim terms [31]. Gene pairs from two different species were considered as positive labels if they share at least one GO term in our selected slim set and negative otherwise.

Hyperparameters that maximized AUPRC between the ranked score matrix and the label generated by our selected slim terms were chosen via 5-fold cross-validation. Importantly, though our GO-based gold standard is based on gene pairs, folds were split by genes, to ensure that the method is only evaluated on genes that it has never “seen.”

We also wanted to evaluate our method on the full set of GO terms (without being restricted to a slim set). To do so, we calculated the Jaccard index between every pair of proteins, i.e., the fraction of GO terms annotated to both proteins in relation to the total number of GO terms annotated to either protein.

### 2.4 Predicting genetic interactions

Genetic interactions for *S. cerevisiae* and *S. pombe* were downloaded from BioGRID (v3.5.187) [28]. Gene pairs with reported “Synthetic Lethality” or “Synthetic Sickness” were regarded as positive examples of Synthetic Sickness and Lethality (SSL). For each species, an equal number of negative examples were subsampled from pairs where both genes were present in the SSL dataset but not reported to show a genetic interaction. Using this gold standard, we applied a support vector machine (SVM) [32] with a radial basis function (rbf) kernel. For a gene pair, the sum of the two corresponding embedding vectors was used as input. As in the cross-validation with GO, folds were split by gene (instead of by gene pair).

### 2.5 Cross-species gene set mapping

For each GO term in the slim set, annotated genes across the two species were considered as matched sets. We calculated a t-score for how closely annotated genes in one species were connected with annotated genes in the other species, while correcting for background network connectivity as in [31]. The final z-score was calculated based on a comparison against a null distribution of gene sets matching in size and degree distribution to each GO term (sampled 100 times).

### 2.6 Clustering cross-species modules

The top 1% of all pairs in the score matrix were used to construct an unweighted and undirected graph between human and mouse, where edges are cross-species alignments between genes. We then applied Louvain [33] to cluster the network vertices and visualized these clusters using Gephi [34] with the OpenOrd [35] layout algorithm (cut parameter = 0.6). Vertices with degrees smaller than 20 were omitted from the final network. For each cluster, we calculated enrichment of GO terms (human and mouse), human OMIM [36] and GWAS [37] disease gene sets, and known human drug targets from DrugBank [38] using the hypergeometric test. All resulting p-values were corrected for multiple hypothesis testing using Benjamini-Hochberg [39].

### 2.7 Existing network alignment methods

We compared our method against three existing gene/protein network alignment methods, MUNK [15], HubAlign [13], and IsoRank [7]. Since both MUNK and ETNA aim to create a joint functional embedding for proteins in two species, we also applied 5-fold cross-validation using the GO functional standard. For HubAlign and IsoRank, due to running time limitations, default parameters were used. We tried running the other network alignment methods that do not use GO as input (for cross-species anchors, Table 1) but were unable to run PrimAlign [8] and L-GRAAL [14]. We were also unable to find a way to give sequence similarity as input to SANA’s [12] code implementation and thus have also excluded it from the comparisons.

### 2.8 Data and code availability

ETNA and all evaluation code was implemented in Python and is available on GitHub (https://github.com/ylaboratory/ETNA), released under the BSD 3-clause license for open source use. BioGRID PPI networks and OrthoMCL orthologs for *S. cerevisiae* and *S. pombe* are included in the data folder as a sample use case.

## 3 Results

### 3.1 Cross-species alignment improves predictive performance of individual embeddings

To identify a joint network embedding, ETNA iterates between calculating individual embeddings and performing cross-training to align the two spaces. Since the joint embedding is only meaningful if functional information captured in the individual network embeddings is preserved, we compared the extent to which the individual network embeddings captured functional signal in the original network, with and without cross-training (i.e., only step 1 of the ETNA framework (Figure 1)). We also checked the performance of still including cross-training, but directly using the adjacency matrix as input to the autoencoders instead of the NetMF matrix *M* (eq. (3)). In each paradigm, we calculated the cosine similarity of latent embeddings for all gene pairs within one species. We evaluated these similarity scores for their ability to recapitulate shared Gene Ontology (GO) annotations between genes within the respective species.

Interestingly, we discovered that the alignment process not only preserves the existing functional relationship captured by the individual network embeddings, but also further improves the predictive performance of the individual embeddings using information from the other network (Figure 2). This suggests that while finding an alignment between two spaces, ETNA’s cross-training step also enables each individual network to leverage the information in the other network to refine its own embeddings. This intuition may also be why cross-training yields a larger performance improvement for *S. pombe* than for *S. cerevisiae*, since *S. cerevisiae* has a much more complete PPI network. We observe a similar trend when comparing ETNA with the modified version that uses the adjacency matrix (instead of the NetMF matrix) as input. The NetMF matrix is able to capture more distant relationships between genes, beyond simply that of the direct neighbors, and thus improves the performance of the network embeddings dramatically.

**Figure 2.**
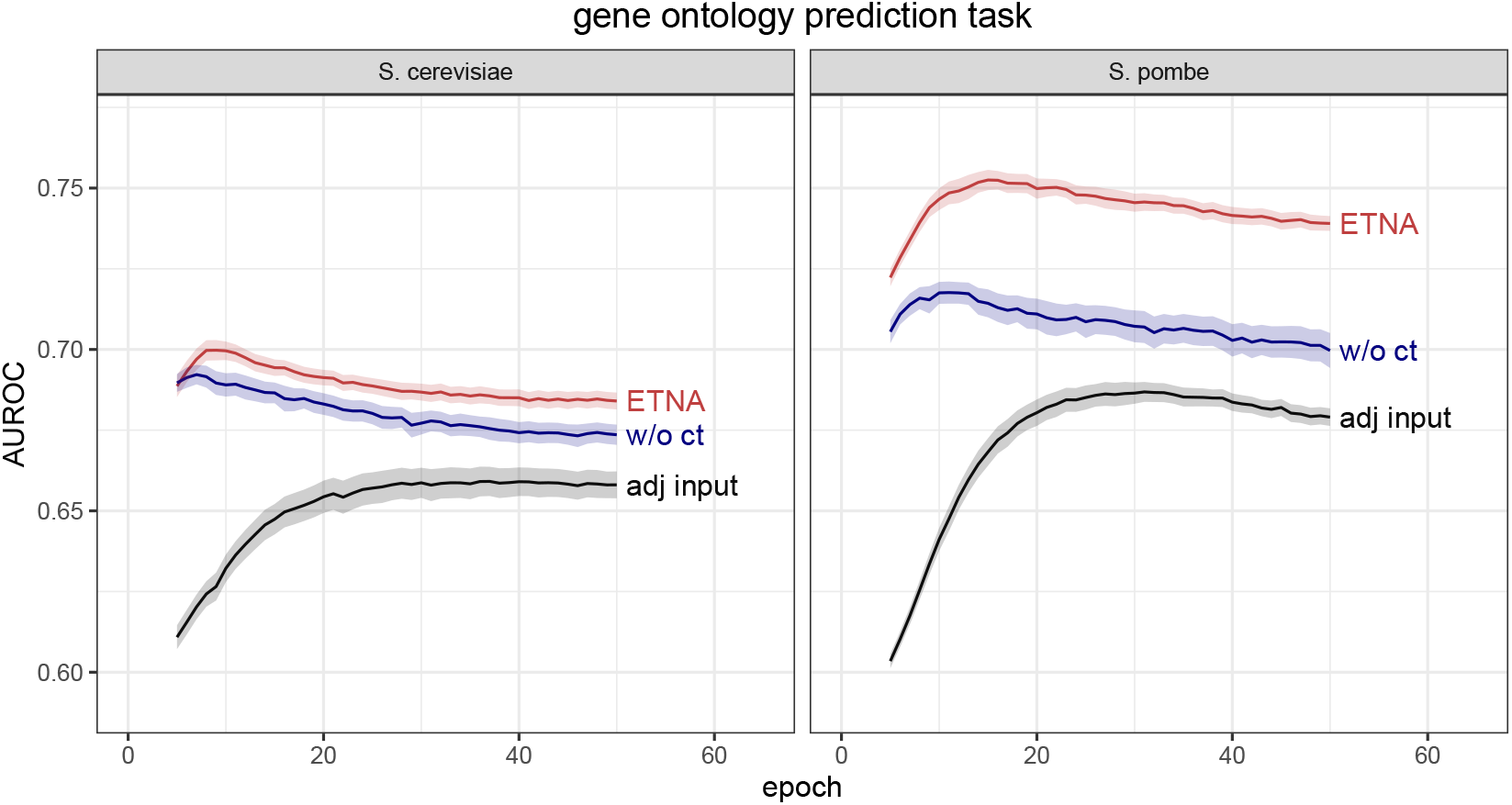
Predicting functional similarity using individual network embeddings in *S. cerevisiae* and *S. pombe* with ETNA (red), embeddings without cross-training (blue), and using the adjacency matrix as input (black). Lines show the mean performance based on 100 random sets of hyperparameters and ribbons denote the 95% confidence interval for predicting functional similarity (defined based on co-annotation to the same GO term (Section 2.3)). Cross-training and using the NetMF matrix instead of the adjacency matrix as input both lead to significant improvements to ETNA’s predictive performance, especially for *S. pombe*. AUROC is shown, and the same trend holds for AUPRC.

### 3.2 ETNA captures cross-organism functional similarity with no prior functional information

We evaluated ETNA’s ability to capture functional similarity given only protein-protein interaction data and sequence-based orthologous gene pairs. Since the majority of functional knowledge transfer tasks involve starting or ending with a human phenotype, we ran ETNA to identify joint embeddings between genes between human and four of the major model organisms: *M. musculus*, *S. cerevisiae*, *D. melanogaster*, and *C. elegans*. For evaluation, we took advantage of the fact that genes across species have been annotated to the same GO terms.

More specifically, to evaluate whether ETNA’s joint embedding captures functionally related gene pairs across species, we examined the predictive performance of the pairwise similarity score matrix (step 3 in Figure 1) against a gold standard based on co-annotation of genes to the same GO term (Section 2.3). We compared ETNA with the predictions made by MUNK, since these are the only two methods that have a joint embedding that outputs a score matrix for pairwise similarities between all genes. Unlike ETNA, MUNK’s predictions are directed from a source organism to a target organism, so we compared ETNA with both of MUNK’s directions.

ETNA generates a more functionally accurate embedding in all tested species pairs, especially for organisms with more complete PPI networks (Table 2). *S. cerevisiae* has the most complete PPI network among all model organisms and is where ETNA has the largest performance improvement over MUNK. Both *D. melanogaster* and *C. elegans* have a sparse PPI network (density < 0.2%); the performance drops for both methods, and the difference between ETNA and MUNK is also correspondingly smaller. Together, these results suggest that with only PPI and sequence ortholog information, ETNA can infer the functional information between genes across species.

**Table 2.** AUPRC over random and AUROC of ETNA and MUNK for predicting cross-species gene pairs that share GO annotations based on 5-fold cross validation. AUPRC over random 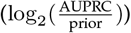 is calculated to facilitate comparisons because each evaluation task has a different prior (ratio of positive examples to negative examples). For the four pairs of species (*H. sapiens*-*M. musculus*, *H. sapiens*-*S. crervisiae*, *H. sapiens*-*D. melanogaster*, *H. sapiens*-*C. elegans*), the random priors are 0.059, 0.043, 0.044, and 0.044 respectively. Because MUNK’s predictions require choosing a source organism and a target organism, we present its performance for both directions (the arrow points from source to target).

| Species Pair | AUPRC (over random) |  | AUROC |  |
| --- | --- | --- | --- | --- |
|  | ETNA | MUNK | ETNA | MUNK |
| *H. sapiens* → *M. musculus* | **0.809** | 0.565 | **0.605** | 0.587 |
| *H. sapiens* ← *M. musculus* |  | 0.445 |  | 0.557 |
| *H. sapiens* → *S. cerevisiae* | **1.376** | 0.810 | **0.677** | 0.627 |
| *H. sapiens* ← *S. cerevisiae* |  | 0.885 |  | 0.626 |
| *H. sapiens* → *D. melanogaster* | **0.706** | 0.602 | 0.603 | 0.597 |
| *H. sapiens* ← *D. melanogaster* |  | 0.619 |  | **0.607** |
| *H. sapiens* → *C. elegans* | **0.622** | 0.512 | **0.596** | 0.572 |
| *H. sapiens* ← *C. elegans* |  | 0.442 |  | 0.554 |

### 3.3 ETNA outperforms existing network alignment and embedding methods in capturing functional similarity

A gene can have multiple functions; thus, the functional similarity between a pair of genes is more complex than the presence or absence of a shared GO annotation. To capture this, we used the Jaccard index to quantify how much functionality is preserved for the cross-species gene pairs. Besides MUNK, two other network alignment methods that are topology-based (IsoRank and HubAlign) are included in the comparison. Besides these existing methods, we also generated two additional baselines: gene pairs ranked by degree and random gene pairs. All of the methods, including ranking by degree, have a higher Jaccard index for higher ranked pairs and gradually converge to random at around 5% of all pairs (data not shown). Node degree is a straightforward but powerful predictor for many network tasks, and a good model should be able to capture information beyond it.

In examining the Jaccard index between the top 5,000 ranked pairs of genes (without orthologs used for alignment) from different methods, we found that ETNA consistently outperforms all other methods across species pairs (Figure 3A). Note that this trend is also consistent across top ranked pairs, with and without orthologs, though we only show 5,000 of them (top ranked pairs) here. In all species pairs except human and worm (*C. elegans*), ETNA is better at capturing gene pairs with multifunctional similarity than existing methods for the top 5,000 pairs. For the comparisons between human and mouse (*M. musculus*), yeast (*S. cerevisiae*), and fly (*D. melanogaster*), ETNA is significantly better than all other methods (one-sided Wilcoxon rank-sum *p <* 10^−16^). For the sparsest PPI network, worm, ETNA has comparable performance to MUNK and is significantly better than the other network alignment methods and baselines (*p <* 10^−6^). This demonstrates that ETNA’s joint embedding can not only reflect whether two genes are related, but also quantify the extent of their relationship.

**Figure 3.**
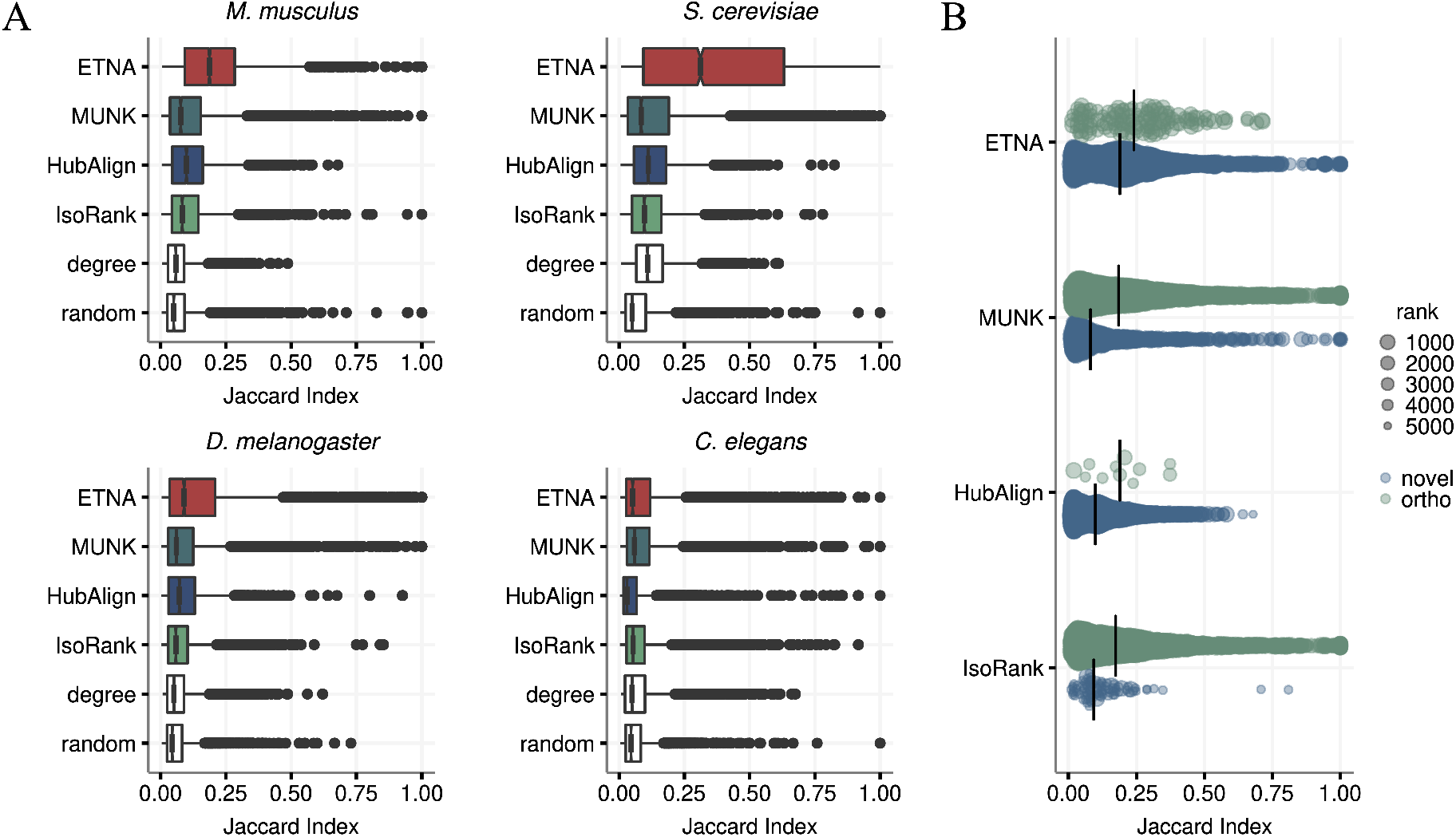
Jaccard index for top 5,000 aligned pairs. (A) Box plot for top 5,000 gene pairs for *H. sapiens* and four model organisms (*M. musculus*, *S .cerevisiae*, *D. melanogaster*, *C. elegans*). Four methods (ETNA, MUNK, HubAlign, IsoRank) and two baselines (degree and random) are compared. ETNA has the best performance in *M. musculus* (*p <* 10^*−*16^), *S. cerevisiae* (*p <* 10^*−*16^), *D. melanogaster* (*p <* 10^*−*16^), and comparable performance to MUNK in *C. elegans*, demonstrating how ETNA’s joint embedding captures multi-functional similarity. All p-values reported are from Wilcoxon rank-sum test [40] to the second best method. (B) Jaccard index comparison between orthologous (ortho) pairs (in blue) and non-orthologous (novel) pairs (in green) within top 5,000 pairs of *H. sapiens*-*M. musculus* alignment. ETNA is better at prioritizing gene pairs beyond orthologs.

When conducting this evaluation, we observed that two of the methods, IsoRank and MUNK, strongly prioritized the orthologs used to anchor the alignment across a species pair (Figure 3B). Unlike other methods, ETNA uses pairwise orthology information alone rather than sequence similarity across all pairs of genes. As IsoRank linearly combines sequence information with topological structures in its algorithm, orthologous pairs will be particularly favored in the alignment. On the other hand, because non-orthologous pairs (most of the pairs) naturally have less sequence similarity information, they are mostly not prioritized. To systematically explore these trends, we also compared each method’s performance with and without orthologous pairs (Figure 3B, human-mouse alignment shown), discovering that ETNA has consistently good performance, beyond prioritizing orthologous pairs alone.

### 3.4 ETNA enables cross-species prediction of genetic interactions

One advantage of having a cross-species joint embedding is that we can detect relationships beyond functional similarity captured by GO. Synthetic sickness and lethality (SSL) is defined as when two (or more) perturbing genes cause a more harmful or deleterious effect on the organism than expected by the combination of single gene perturbations. Synthetic lethality has received increasing attention for its potential application to cancer treatments [3], but the experimental detection of SSL is inefficient and hard to perform, especially on higher order organisms. We first used SSL information from both *S. cerevisiae* (*Sce*) and *S. pombe* (*Spo*) together to predict the held out SSL pairs (Section 2.4). ETNA shows a significant performance increase in both *Sce* and *Spo* in comparison to MUNK (regardless of which directionality was used) (Table 3).

**Table 3.** ETNA consistently outperforms MUNK in prediction of genetic interactions between *S. cerevisiae* (*Sce*) and *S. pombe* (*Spo*). For each SSL prediction task, *A → B* indicates SSL pairs in *A* were used for training to predict SSL pairs in *B* (e.g., *Sce* + *Spo → Sce* indicates that SSL gene pairs from both *Sce* and *Spo* were used for training, and a set of held out SSL gene pairs in *Sce* were used for evaluation). Notably, ETNA is able to predict genetic interactions across species well using *only* interactions reported in the other (*Sce→Spo*, *Spo→Sce*). Each of the gold standards had balanced positives and negatives (prior=0.5).

| SSL Prediction Task | AUPRC |  |  | AUROC |  |  |
| --- | --- | --- | --- | --- | --- | --- |
|  | ETNA | MUNK |  | ETNA | MUNK |  |
| | $Sce \leftrightarrow Spo$ | $Sce \rightarrow Spo$ | $Sce \leftarrow Spo$ | $Sce \leftrightarrow Spo$ | $Sce \rightarrow Spo$ | $Sce \leftarrow Spo$ |
| $Sce + Spo \rightarrow Sce$ | **0.798** | 0.634 | 0.694 | **0.768** | 0.622 | 0.680 |
| $Sce + Spo \rightarrow Spo$ | **0.772** | 0.662 | 0.648 | **0.776** | 0.668 | 0.669 |
| $Sce \rightarrow Spo$ | **0.730** | 0.611 | 0.617 | **0.740** | 0.613 | 0.614 |
| $Spo \rightarrow Sce$ | **0.708** | 0.612 | 0.565 | **0.681** | 0.563 | 0.446 |

Next, we tackled the more challenging task of only using genetic interactions reported in one yeast species to predict the SSL pairs in the other. This task more closely mimics how we may use these embeddings in practice, where specific types of data may only be richly available in one model organism. We evaluate how well we can “transfer” the wealth of genetic interactions captured in *Sce* to other species using only PPI and sequence information, which are more readily available across species. As expected, the predictive performance for this challenging task is worse than using SSL information from both species as input. Nevertheless, ETNA still performs very well and also significantly better than MUNK. Notably, we also noticed MUNK’s joint embeddings can result in inconsistent predictions between the two alignment directions, and furthermore, sometimes the non-intuitive direction may perform better for a particular task. For example, for the task of predicting genetic interactions for *Sce* using only *Spo* gene pairs (*Spo* → *Sce*), MUNK’s *Sce* → *Spo* performs better whereas the matching direction has near random performance. This highlights the importance of having a bidirectional embedding, and in general, these results demonstrate how ETNA enables the knowledge transfer of genetic interactions from a well-studied species to another.

### 3.5 ETNA’s joint embedding space captures functional similarity of gene sets

Researchers also often encounter gene sets of interest, and as many important biological processes are performed by the cooperation of multiple proteins, beyond matching gene pairs between two species, it is important to explore the functional alignment of gene sets. We thus explored whether genes annotated to the same GO term across two species were more significantly connected to each other than expected by random.

To this end, as in Greene et al [31], we compared the connectivity of the GO term split across species to a null distribution of random degree-matched gene sets of the same size (Section 2.5). Across the board, ETNA was able to identify significant (after Bonferroni correction [41, 42]) matches for over 75% of the GO terms shared between species (*H. sapiens*-*S. cerevisiae*: 94%, *H. sapiens*-*M. musculus* 75%, *H. sapiens*-*D. melanogaster* 77%, and *H. sapiens*-*C. elegans* 82%), whereas other methods were only able to match 25% of the GO terms at the same significance threshold (Figure 4). In this evaluation, we show that ETNA not only captures pairwise gene relationships between two species but also that functional groups are also meaningfully clustered in the joint embedding space.

**Figure 4.**
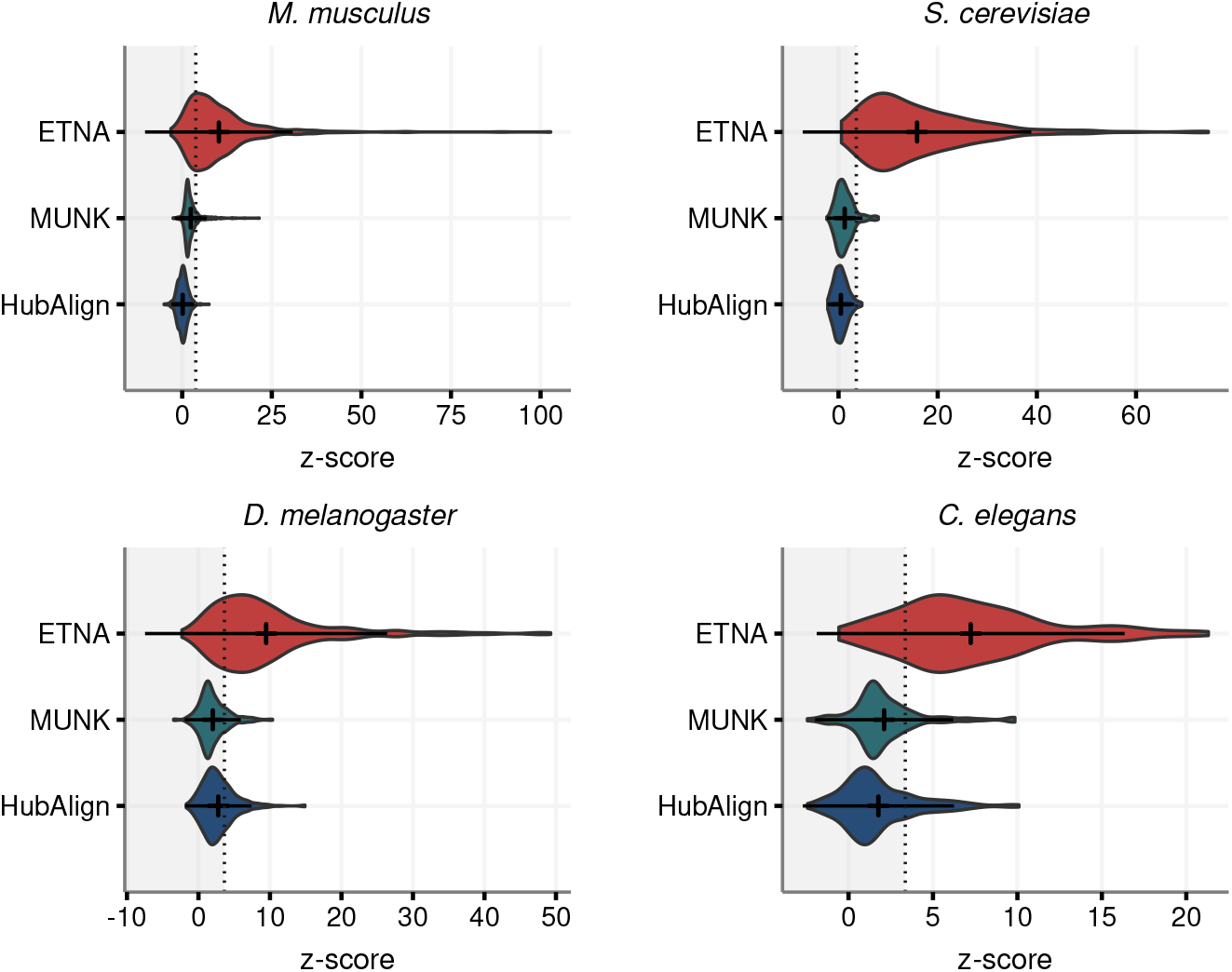
Evaluating cross-species GO term matching demonstrates that functional gene sets have consistently stronger correspondence than other methods. The alignment between *H. sapiens* and four model organisms (*M. musculus*, *S. cerevisiae*, *D. melanogaster*, *C. elegans*) are considered. There are 645, 292, 349, 129 shared GO terms, respectively. We calculated a z-score for connectivity between genes annotated to the same GO term across species against a null distribution generated from random degree-matched gene sets of the same size. The dotted line is the z-score that corresopnds to the Bonferroni corrected p-value. We were unable to evaluate IsoRank for this task because it only outputs the top 100,000 pairs and not a full list of predictions.

By taking a closer look at these z-scores, we found that most of the high scoring GO terms described functions such as RNA polymerase, transcription, etc. These functions have been shown that are highly conserved through evolution across eukaryotic organisms. As for GO terms that had a z-score below the Bonferroni threshold, many were child nodes under the umbrella GO term “response to stimulus.” One hypothesis for why ETNA does not capture these functions as well is that stimulus response may not involve as many physically interacting genes (e.g., via signal transduction, phosphorylation), so when only considering the PPI network as input, ETNA could miss these relationships.

### 3.6 ETNA can reveal shared mechanisms of fundamental biological processes and disease pathology

So far, our evaluations have shown that ETNA captures known functional biological processes. Taking a closer look at the alignment between human and mouse, we wanted to explore whether the top gene pairings can be used for functional knowledge transfer of other gene sets. Thus, we took the top 1% of ETNA scores between human and mouse and clustered the data, identifying functional modules (Section 2.6). We found 122 modules with at least 5 genes and performed enrichment on GO terms (human and mouse), drug targets (DrugBank [38]), and known human disease genes (annotated in OMIM [36]). There were a range of significantly enriched terms in most modules, but here we highlight the top 10. These modules covered a range of interesting disease mechanisms and key conserved core biological processes, including mitochondrial processes, developmental signal, and extracellular matrix-associated processes (Figure 5).

**Figure 5.**
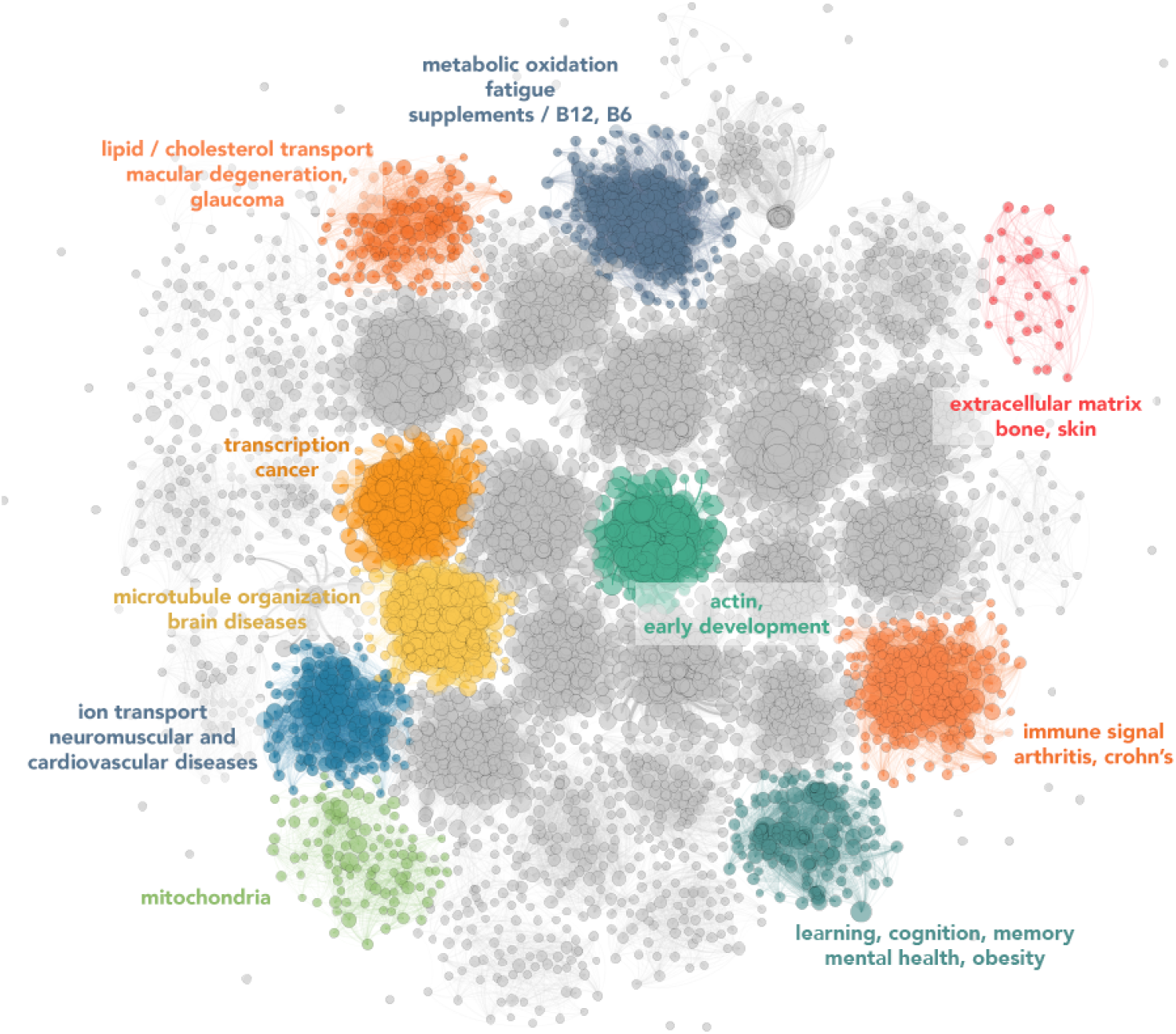
Network of the top gene mappings between human and mouse. This network shows the top 1% of ETNA scores for the alignment of *H. sapiens* and *M. musculus*, where each vertex is a gene and each edge is an alignment between the two species. Enrichment of disease, drug, and GO terms were calculated for each cluster, and the top 10 most enriched clusters are colored and labeled here with significant terms summarized, highlighting the potential for ETNA to discover novel biology.

An interesting module (orange, middle left) was enriched for GO terms in the human and mouse pertaining to proliferation, transcription, and apoptosis, and cancer related OMIM genes of the breast, reproductive tract, and the gastrointestinal system, suggesting similarities between these classes of cancer types. The lipid/cholesterol transport cluster (orange, upper left) interestingly also has a significant enrichment for several eye-related diseases, and it has been well-documented that high cholesterol levels have been shown to increase the risk of both macular degeneration and glaucoma [43, 44, 45, 46]. Furthermore, the learning, cognition, and memory cluster (teal, bottom right) also captures cross-connections between mental health disorders as well as well-supported links to gut and obesity [47, 48, 49]. Several relevant drug targets were also enriched in this cognition cluster, with many having sedative and/or anti-anxiety indications (e.g., clonazepam, flurazepam, oxazepam, etc.). Notably, weight gain is also a side effect of many of these “-zepam” drugs, suggesting ETNA may be uncovering a shared mechanism between obesity and processes in the brain. While we highlight a few specific examples here, there are many more interesting connections that are ripe for exploration, and as such all models, embeddings, scores, and code will be made available for download.

## 4 Discussion

In this study, we introduce ETNA as a method to transfer functional information across species. Unlike traditional network alignment methods that calculate a single score to capture similarity between a pair of genes, ETNA generates a general purpose joint embedding, capturing functional relevance between two genes as multi-dimensional vectors. Instead of linearly combining topological structure and orthologous information, ETNA introduces an autoencoder-based framework that captures the non-linearities, as well as local and global relationships in network topology, then uses cross-training to construct a joint embedding.

The evaluations have shown that ETNA is capable of capturing both pairwise and group functional relationships between human and other model organisms. Beyond inferring unannotated gene functions from their closely related genes in other organisms, ETNA’s embedding enables transfer of genetic interaction knowledge from one species to another. As the number of possible pairs of genetic interactions has a combinatorial relationship with the number of genes, and gene knockout can be costly or intractable to perform at scale on higher-order organisms, ETNA’s joint embedding provides a new way to unravel gene relationships that are difficult to detect experimentally. Finally, by exploring the human-mouse functional landscape, we are able to identify interesting connections between mouse functional studies with complex human diseases, setting the stage for potential opportunities for translational studies.

Though we have applied ETNA to PPI networks here, the methodological framework can be easily applied to other types of biological networks. As shown in Section 3.2, the more complete a PPI is, the better ETNA can use this information to create a more accurate joint embedding, so it would be interesting to explore whether using predicted PPIs to supplement experimentally derived networks would improve performance. But beyond PPI networks, we can envision alignment of metabolic or regulatory networks. Integrated functional networks [50] designed to predict functional similarities would also be natural to use as input into ETNA.

Furthermore, because the joint embedding of ETNA does not require choosing a source and target, it opens possibilities for extending the framework to simultaneously perform alignment for more than two species. We have found here that the cross-training step enables ETNA to use information from other species to refine individual embeddings, so we anticipate that a “multiple-species network alignment” could result in an even more accurate joint embedding and enable placing model systems with limited experimental studies into the functional landscape.

## Notes

### Competing Interest Statement

The authors have declared no competing interest.

